# Cryptic and extensive hybridization between ancient lineages of American crows

**DOI:** 10.1101/491654

**Authors:** David L. Slager, Kevin L. Epperly, Renee R. Ha, Sievert Rohwer, Chris Wood, Caroline Van Hemert, John Klicka

## Abstract

Most species and therefore most hybrid zones have historically been described using phenotypic characters. However, both speciation and hybridization can occur with negligible morphological differentiation. The Northwestern Crow (*Corvus caurinus*) and American Crow (*Corvus brachyrhynchos*) are sister taxonomic species with a continuous distribution that lack reliable traditional characters for identification. In this first population genomic study of Northwestern and American crows, we use genomic SNPs (nuDNA) and mtDNA to investigate whether these crows are genetically differentiated and the extent to which they may hybridize. We found that American and Northwestern crows have distinct evolutionary histories, supported by two nuDNA ancestry clusters and two 1.1%-divergent mtDNA clades dating to the late Pleistocene, when glacial advances may have isolated crow populations in separate refugia. We document extensive hybridization, with geographic overlap of mtDNA clades and admixture of nuDNA across >1,400 km of western Washington and western British Columbia. This broad hybrid zone consists of late-generation hybrids and backcrosses, not recent (e.g., F1) hybrids. Nuclear DNA and mtDNA clines were both centered in southwestern British Columbia, farther north than previously postulated. The mtDNA cline was narrower than the nuDNA cline, consistent with Haldane’s rule but not sex-biased dispersal. Overall, our results suggest a history of reticulate evolution in American and Northwestern crows, consistent with potentially recurring neutral expansion(s) from Pleistocene glacial refugia followed by lineage fusion(s). However, we do not rule out a contributing role for more recent potential drivers of hybridization, such as expansion into human-modified habitats.

## Introduction

Phenotypic characters have historically been a primary basis for distinguishing between species (Bickford et al. 2007), but the increasing availability of DNA evidence has made clear that speciation is not always accompanied by morphological change (Fiser et al. 2018). Genomic analyses of morphologically conserved groups have uncovered cryptic species across the tree of life (e.g., Pfenninger and Schwenk 2007, Satler et al. 2013, Hotaling et al. 2016, Larsen et al. 2017), yet not all morphologically diagnosed species show genomic differences (e.g., Mason and Taylor 2015).

Morphological similarity between closely related species sometimes reflects divergence too recent for morphological change to have occurred (Egea et al. 2016). For species with a longer history of evolving independently, morphological similarity not due to convergence may reflect a lack of directional selection on visually salient morphological traits (Bickford et al. 2007) or stabilizing selection for a common ancestral morphology (i.e., morphological stasis, e.g., Smith et al. 2011). Speciation without morphological differentiation may be especially prevalent in morphologically austere taxa (e.g., sponges, Klautau et al. 1999), where visual cues are less important for species recognition or mating signals, or in taxa undergoing morphological stasis (Bickford et al. 2007). In addition, some apparent cases of cryptic speciation may simply reflect taxonomic neglect of the relevant morphological characters rather than a true lack of morphological change (e.g., in coccolithophores, Saez et al. 2003).

Hybrid zones are classic and powerful natural laboratories for understanding the various stages of the speciation continuum (Harrison 1993, de Queiroz 1998). Geographic patterns at hybrid zones can illuminate processes like the evolution of reproductive isolation, the influence of differing levels of gene flow, and the fusion of lineages that can occur when previously allopatric populations experience secondary contact. However, because most species have historically been diagnosed morphologically, most hybrid zone studies have likewise focused on morphologically distinct parental species (Barton and Hewitt 1989). By comparison, research addressing morphologically cryptic hybrid zones is less common and has begun to appear only recently (e.g., Pfenninger and Nowak 2008, Herrera-Aguilar et al. 2009, Patel et al. 2015, Quilodran et al. 2018, Pulido-Santacruz et al. 2018). These few studies have already documented a variety of evolutionary patterns and processes operating in the absence of morphological differences, including cryptic reticulate evolution (Kearns et al. 2018). However, additional genomic studies of cryptic hybrid zones are needed to better synthesize how evolutionary processes may differ at hybrid zones with and without morphological differentiation.

The Northwestern Crow (*Corvus caurinus*) and American Crow (*Corvus brachyrhynchos*) are sister taxa (Haring et al. 2012) that have long been considered separate species (Baird 1858). However, they are nearly identical morphologically, and collectively they have a continuous distribution along the Pacific coast of North America (Figure 1; Clements et al. 2017). The traditional, phenotypic characters for identifying these all-black corvids are based on putative differences in morphology, voice, and ecology, and have always been controversial (see Discussion). Uncertainty in identification of these crows has moreover led to widely varying interpretations of the precise location of their range boundaries and contention regarding the nature and extent of a secondary contact zone. Interpretations of the latter have ranged from assortative mating of discrete forms in sympatry (Brooks 1917, 1942) to clinal variation without diagnosable differences (Johnston 1961), and little new information has surfaced during the past half century. The only prior molecular study sequenced mtDNA from just a handful of American and putative Northwestern crows, and all three “Northwestern” Crow samples were from near the range boundary, leaving doubts as to whether these were Northwestern Crows, American Crows, or hybrids (Haring et al. 2012).

**Figure 1:**
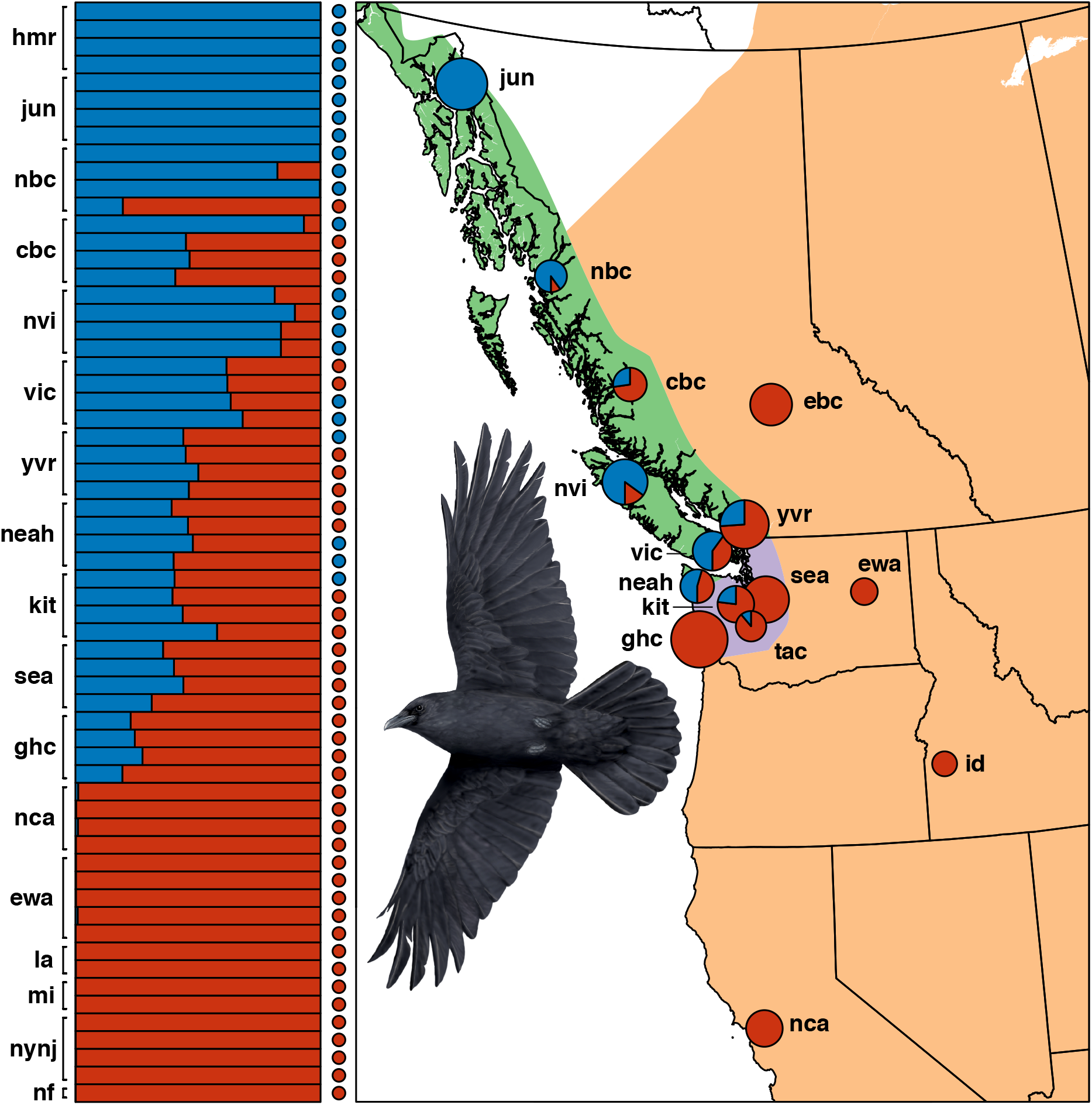
Extent of hybridization between Northwestern Crow (blue) and American Crow (red). At left, bars show nuDNA K=2 ancestry proportions and adjacent circles indicate mtDNA haplogroup for the same 62 individuals. At right, locality pies depict mtDNA haplogroup proportions from the full mtDNA dataset (n=6-31 per locality), and background colors indicate range maps for Northwestern Crow (green), American Crow (orange), and the overlap zone (purple; BirdLife 2013). Sample localities outside the mapped region contained 100% Northwestern or 100% American mtDNA haplogroups (Table S1, Figure S2). Sample IDs for bars at left run numerically from top to bottom (e.g., *mi01, mi02*; see Table S1). Original crow illustration by Kevin L. Epperly.

In this study, our objective was to use genomic data and geographically robust sampling of American and Northwestern crows to better understand the evolutionary history of these presumptive species that lack well-defined phenotypic characters. Specifically, we set out to 1) assess whether Northwestern and American crows represent independently evolving evolutionary lineages, and, if so, 2) determine the extent to which they might hybridize or exhibit reproductive isolation, and 3) better understand their evolutionary and biogeographic history.

## Materials and Methods

### Sample collection and DNA extraction

To conduct a population genetic survey of American Crows and Northwestern Crows near their range boundary and across North America, we sampled tissue (n=218), blood (Alaska; n=35), or feather material (Idaho; n=6) from crows identified *a priori* as either species (Table S1, Figure S1). We also included two Carrion Crow (*Corvus corone*) tissue samples as outgroups (Haring et al. 2012). Tissue samples were obtained from natural history museums and were generally associated with vouchered specimens (Table S1). Blood samples were obtained under permits and approvals from the US Fish and Wildlife Service, the Alaska Department of Fish and Game, and the Institutional Animal Care and Use Committees at the University of Alaska Fairbanks and the US Geological Survey Alaska Science Center. Feather samples were collected under permits from the US Fish and Wildlife Service and the Idaho Department of Fish and Game, following recommended protocols in the *Guidelines to the Use of Wild Birds in Research* (Gaunt et al. 1997). We extracted total genomic DNA with a DNeasy tissue extraction kit (Qiagen, Valencia, CA) following manufacturer protocol.

### Mitochondrial DNA (mtDNA) sequencing

To survey a large sample of crows across the putative contact zone and throughout North America and to conduct divergence dating, we amplified 1,041 base pairs (bp) of mtDNA NADH dehydrogenase subunit 2 (ND2) from 259 individuals (Table S1). We used primers L5215 (Hackett 1996) and TrC (Miller et al. 2007) and 12.5 *μ*L PCR reactions on a T100 thermal cycler (Bio-Rad, Hercules, CA) as follows: 94°C for 2.5 min, 35 cycles of 94°C for 30 s, 54°C annealing for 30 s, 72°C for 1 min), 10 min at 72°C, 10°C hold. We sent PCR products to the High-Throughput Genomics Unit at the University of Washington for cleanup and sequencing. We unambiguously aligned complementary strands with Sequencher 5.0 (Gene Codes Corporation, Ann Arbor, MI) and downloaded a Carrion Crow ND2 sequence from GenBank as an outgroup (Table S1).

### mtDNA haplotype network and divergence dating

We constructed a median-joining ND2 haplotype network (Bandelt et al. 1999) in PopArt 1.7 (Leigh and Bryant 2015) and calculated mean pairwise distance between haplotype groups in R (R Core Team 2016) using ape 3.5 (Paradis et al. 2004). We omitted one sample (gws4003_Pierce_Co) from the haplotype network and pairwise divergence calculations because only a partial ND2 sequence was obtained. We estimated divergence dates using an ND2 sequence evolution rate of 2.9×10^−2^ substitutions site^−1^My^−1^ (95% HPD interval 2.4 − 3.3 ×10^−2^ derived from Hawaiian honeycreepers (Drepanidinae) and calibrated using sequential uplift dates of Hawaiian islands (Lerner et al. 2011).

### Nuclear DNA (nuDNA) SNP library preparation

To assess whether Northwestern and American crows show distinct evolutionary histories in the nuclear genome and to determine the extent to which these crows hybridize, we generated reduced representation double digest restriction-associated DNA (ddRAD) SNP libraries. We generated these genomic SNP libraries from a subset of 62 American/Northwestern Crow individuals for which we had also sequenced mtDNA ND2, and two Carrion Crows as an outgroup. To maximize our power to detect potentially distinct, sympatric lineages, we used mtDNA results as a guide for selecting individuals for nuDNA analysis. At localities containing both of the major mtDNA ND2 haplogroups in our broad population genetic survey, we selected individuals for ddRAD sampling to reflect the approximate overall ratio of mtDNA haplogroups at each locality. We followed the ddRAD sequencing protocol of Peterson et al. (2012). We verified high molecular weight DNA on a gel and quantified DNA using a Qubit 2.0 Fluorometer (Invitrogen, Carlsbad, CA) and a Qubit dsDNA High Sensitivity Assay Kit (Thermo Fisher Scientific, Waltham, MA). We digested 350-500 ng of DNA in a 50*μ*L reaction in CutSmart Buffer with SbfI-HF and MspI restriction endonucleases (New England BioLabs, Ipswich, MA). We verified digestion by eye on a gel. After a bead cleanup (Rohland and Reich 2012), we quantified DNA and diluted samples to the concentration of the least concentrated sample (≥10 ng/*μ*L). We ligated P1 adapters (8 unique adapters per pool of 8, differing by at least 2 bp) to SbfI ends and P2 adapters to MspI ends at 4.5x fold excess with T4 DNA Ligase (New England BioLabs, Ipswich, MA) in 40*μ*L reaction volumes, incubating at 37°C for 30 min and 65°C for 10 min. We pooled each set of 8 samples and re-quantified DNA after two bead cleanups. We size-selected 415-515 bp DNA fragments on a BluePippin machine (Sage Science, Beverly, MA) and re-quantified DNA. We performed a 50-*μ*L PCR reaction using Phusion High-Fidelity DNA Polymerase (New England BioLabs, Ipswich, MA), PCR primer 1, and a uniquely indexed PCR primer 2 for each pool of 8 (Peterson et al. 2012). We performed a bead cleanup after PCR and re-quantified DNA with the Qubit 2.0 fluorometer. We then quantified DNA fragment peak size and molarity using an Agilent 2200 TapeStation (Agilent Technologies, Santa Clara, CA). We multiplexed 96 avian ddRAD libraries from this study and another study, obtaining two runs of single-end 50-bp reads from Illumina HiSeq 2500 at the Computational Genomics Research Laboratory at the University of California (Berkeley, CA).

### nuDNA sequence assembly

We used *process_radtags* in STACKS 1.42 (Catchen et al. 2013) with default settings to demultiplex reads, discard low-quality reads, discard reads with an uncalled base, rescue barcodes, and rescue SbfI RAD tags. To align reads to an American Crow reference genome (Zhang et al. 2014), we built a genome database using default settings for *gmap_build* in GMAP (Wu and Watanabe 2005). We aligned reads to the reference genome with GSNAP version 2016-09-23 (Wu and Nacu 2010), specifying ≥ 90% coverage, ≤ 3 mismatches, and default settings for other parameters. We converted alignments to BAM format with SAMtools (Li et al. 2009) and created loci and called SNPs from the aligned reads with *ref_map.pl* in STACKS 1.42 (Catchen et al. 2013). We constructed a 64-crow alignment including the two Carrion Crow outgroup samples and a 62-crow alignment containing only American and Northwestern crows. For each dataset, we retained stacks with a depth of ≥ 3 identical reads, loci with a coverage of ≥ 4 samples, and bi-allelic SNPs with a minor allele count ≥ 2 and heterozygosity ≤ 0.5. We output Structure files for downstream analysis. We used a custom R script to generate 64-crow and 62-crow alignments of unlinked SNPs retaining one random SNP per locus, a 62-crow alignment without missing data, and 48-crow alignments of Pacific coastal birds with and without missing data.

### nuDNA cluster analyses

We used the K-means clustering algorithm implemented in the *find.clusters* function in Adegenet 2.0.1 (Jombart et al. 2010) to calculate the Bayesian information criterion (BIC) for K=1 through K=10. We used the 62-sample dataset with no missing data after transforming by principal components analysis (PCA), retaining all principal components.

We also used the Bayesian admixture model with correlated allele frequencies implemented in Structure 2.3.4 (Pritchard et al. 2000) to conduct 5 replicate runs each for K=1 to K=3 with 200,000 generations and a burn-in of 20,000. We estimated the number of clusters in the unlinked 62-sample dataset with missing data by analyzing the rate of change in the likelihood distribution between successive K values in Structure Harvester 0.6.94 (Evanno et al. 2005, Earl and vonHoldt 2012). We combined results from replicate runs while accounting for permutations and label switching using CLUMPP 1.1.2 (Jakobsson and Rosenberg 2007).

### Recent-generation vs. late-generation hybrids

We compared genomic hybrid indices to inter-taxon heterozygosities to determine whether individual crows were recent-generation hybrids or descendants of long-admixed populations (Milne and Abbott 2008, Bouchemousse et al. 2016). We designated crows as parental American or parental Northwestern if these respective ancestry proportions exceeded 0.98 in our combined K=2 Bayesian clustering runs (Scordato et al. 2017). We used a custom R script to subset the 62-sample SNP alignment containing missing data to include only parental individuals and only variable, unlinked SNPs present for ≥75% of individuals. With this alignment we calculated a SNP-specific F_*ST*_ between parental populations with R package Hierfstat (Goudet 2005). We considered SNPs with F_*ST*_ > 0.6 to be ancestry-informative (Scordato et al. 2017) and further limited our alignment to these SNPs using a custom R script. For each individual crow, we calculated a maximum likelihood estimate of the genomic hybrid index and the average inter-taxon heterozygosity across ancestry-informative loci using R package Introgress (Gompert and Buerkle 2010). F1 hybrids have an expected hybrid index of 0.5 and expected heterozygosity of 1.0 for loci fixed in parental individuals. Heterozygosity is reduced in later-generation hybrids and backcrosses. We considered crows with hybrid index >0.25 and <0.75 and heterozygosity >0.5 to be recent-generation hybrids, individuals with hybrid index >0.25 and <0.75 but with heterozygosity <0.5 to be later-generation hybrids, and birds with hybrid index <0.25 or >0.75 to be backcrosses (Milne and Abbott 2008, Larson et al. 2014, Scordato et al. 2017, Toews et al. 2018).

### Fitting and comparing clines for mtDNA and nuDNA

We fit cline models for mtDNA and nuDNA along the Pacific Coast using R package Hzar (Derryberry et al. 2014). We assigned transect distances to each Pacific coastal population using a smoothed curve drawn parallel to the Pacific coastline in ArcMap (ESRI 2011). For mtDNA, we fit a cline model for population frequencies of American mtDNA (range 0 to 1) using *hzar.doMolecularData1DPops* and *hzar.makeCline1DFreq.* For nuDNA, we used an alignment with no missing data for 48 crows along the Pacific coastal transect. We extracted scores for the first principal components analysis (PCA) axis (PC1) of this alignment in Adegenet (Jombart 2008). To match the scale used for mtDNA haplogroup frequencies, we applied a custom linear transformation to nuDNA PC1 such that our southernmost and northernmost localities had population means of 0 and 1, respectively. We fit a cline model to genomic population means and variances using *hzar.doNormalData1DPops* and *hzar.make Cline lD Normal.* For both mtDNA and nuDNA, we fit models without exponential tails and restricted parameter search space to a liberal yet reasonable range of values (cline center >1000 km and <6000 km; cline width <7000 km; Derryberry et al. 2014). We ran the Markov chain Monte Carlo (MCMC) optimizer for 3 iterative cycles using *hzar.chain.doSeq*, retaining the third run for subsequent analysis (Derryberry et al. 2014). We used *hzar.get.ML.cline* and *hzar.getLLCutParam* to obtain maximum likelihood estimates for cline fits, cline centers ± 2 log likelihood (LL), and cline widths ± 2 LL.

### Assessing heterogeneity of admixture near geographic features of interest

To provide an ad hoc assessment of potential differences in gene flow and admixture between Vancouver Island crows and those across the water barrier on the adjacent mainland, we compared nuDNA ancestry proportions of individuals from Vancouver Island (*vic* + *nvi* localities, n=8) to those of nearby mainland populations (*yvr* + *cbc* localities, n=8). We compared Northwestern nuDNA ancestry from the combined K=2 Structure runs using a Student’s t-test with a two-tailed hypothesis and pooled variances. We also compared ND2 haplogroup proportions of crows between Vancouver Island the adjacent mainland using a Pearson’s chi-squared test with 2,000 Monte Carlo replicates (n=35 for *nvi* + *vic*; n=34 for *cbc* + *yvr*).

To assess the Skeena River valley of British Columbia as a potential corridor for gene flow across the Coast Mountains, we conducted an ad hoc comparison of the within-population variance in nuDNA K=2 ancestry proportions between two localities nearest to the Skeena River and all other localities. We fitted two nested mixed effects models in nlme (Pinheiro et al. 2017) and compared them with a likelihood ratio test. Both models included the Northwestern nuDNA ancestry proportion as the response variable, membership in *nbc* + *cbc* (the localities nearest to the Skeena River) as a fixed effect, and locality as a random effect. The more complex model incorporated a variance function parameter allowing the Skeena River localities to take on a common within-population variance estimate that differed from a within-population variance estimate common to all other populations.

## Results

### Sequence alignments

#### nuDNA

We obtained 181,580,191 raw sequencing reads across 64 samples in 8 pools, retaining 150,638,152 reads (83.0%) after discarding 13,626,576 (7.5%) with ambiguous barcodes, 1,988,891 (1.1%) with ambiguous RAD tags, and 15,326,572 (8.4%) with low quality scores. The median coverage depth of samples processed by *refmap.pl* was 118x (range 33x-187x). The Bayesian admixture model readily distinguished the two Carrion Crow samples from the 62 American and Northwestern crows, and we excluded these Carrion Crows from subsequent analyses. The 62-sample alignment contained 7,292 loci with 9,563 SNPs before random sub-sampling to 7,292 unlinked SNPs. The 62-sample alignment with no missing data contained 738 unlinked SNPs, and the 48-sample alignment of Pacific coastal samples with no missing data contained 905 unlinked SNPs.

#### mtDNA

We obtained 259 mtDNA ND2 sequences, including 258 full-length ND2 sequences (1041 bp) and one partial sequence identifiable to haplogroup.

### American and Northwestern crows have distinct evolutionary histories

Genomic SNPs and mtDNA sequences both revealed distinct evolutionary histories for Northwestern and American crows. Two nuDNA ancestry clusters corresponding to Northwestern and American crows (Figure 1) were supported by two different clustering approaches. The K-means algorithm minimized the BIC at K=2 (256.85 for K=1, 255.12 for K=2, and ≥255.81 for 3≤K≤10). The mean log likelihood of the Bayesian admixture model began decelerating at K=2 (−174,866 for K=1, −163,293 for K=2, −161,074 for K=3). Combined results from replicate K=2 Bayesian clustering runs are presented in Figure 1.

A median joining mtDNA ND2 haplotype network also revealed two distinct haplogroups (Figure S1). These haplogroups were separated by 3 fixed differences and a mean uncorrected pairwise distance of 1.10% (range 0.58% to 1.63%), and we hereafter refer to these haplogroups as Northwestern (n=95) and American (n=164). The estimated time of divergence was ~381,000 years ago (95% HPD interval 335-460 kya).

### American and Northwestern Crow mtDNA haplogroups overlap geographically

Overall, population frequencies of the Northwestern mtDNA haplogroup increased with latitude west of the Cascades and Coast Ranges. All Alaskan crows had the Northwestern mtDNA haplogroup, and all crows from California, east of the Cascades of Oregon and Washington, and east of the Coast Mountains of British Columbia had the American mtDNA haplogroup (Figure 1). Individual crows with American and Northwestern mtDNA haplogroups co-occurred within a >1,400 km overlap zone on the Pacific slope of Washington and British Columbia. We found no additional geographic-genetic structuring within the Northwestern or American mtDNA haplogroups.

### American and Northwestern crows hybridize extensively

Of the crows for which we sampled nuDNA, all individuals from Alaska had pure (>98%) Northwestern ancestry, and all from California and east of the Cascades/Coast Mountains had pure American ancestry (Figure 1). Hybrid crows with nuDNA ancestry >2% and <98% occupied a Pacific coastline distance of >1,500 km and included all of the crows that we sampled from coastal Washington to coastal British Columbia at the approximate latitude of northern Haida Gwaii. Among four crows at this northern limit of the hybrid zone, two had pure Northwestern ancestry, one hybrid had mostly Northwestern ancestry, and another hybrid had mostly American ancestry.

### The hybrid zone consists of late-generation hybrids and backcrosses

Among 62 Northwestern and American crows with nuDNA SNP data, 10 were parental Northwestern, 18 were parental American, and 34 were hybrids. Our alignment of the 28 parental crows contained 3,582 unlinked SNPs present in ≥75% of individuals. Thirty-five SNPs were ancestry-informative (F_*ST*_ > 0.6) between parental American and parental Northwestern crows, including 2 SNPs with fixed differences (F_*ST*_ = 1).

All 28 hybrids we sampled had low inter-taxon heterozygosities at ancestry-informative SNPs given their respective hybrid indices, indicating that they were late-generation hybrids and backcrosses and not F1s or early-generation hybrids (Figure 2).

**Figure 2:**
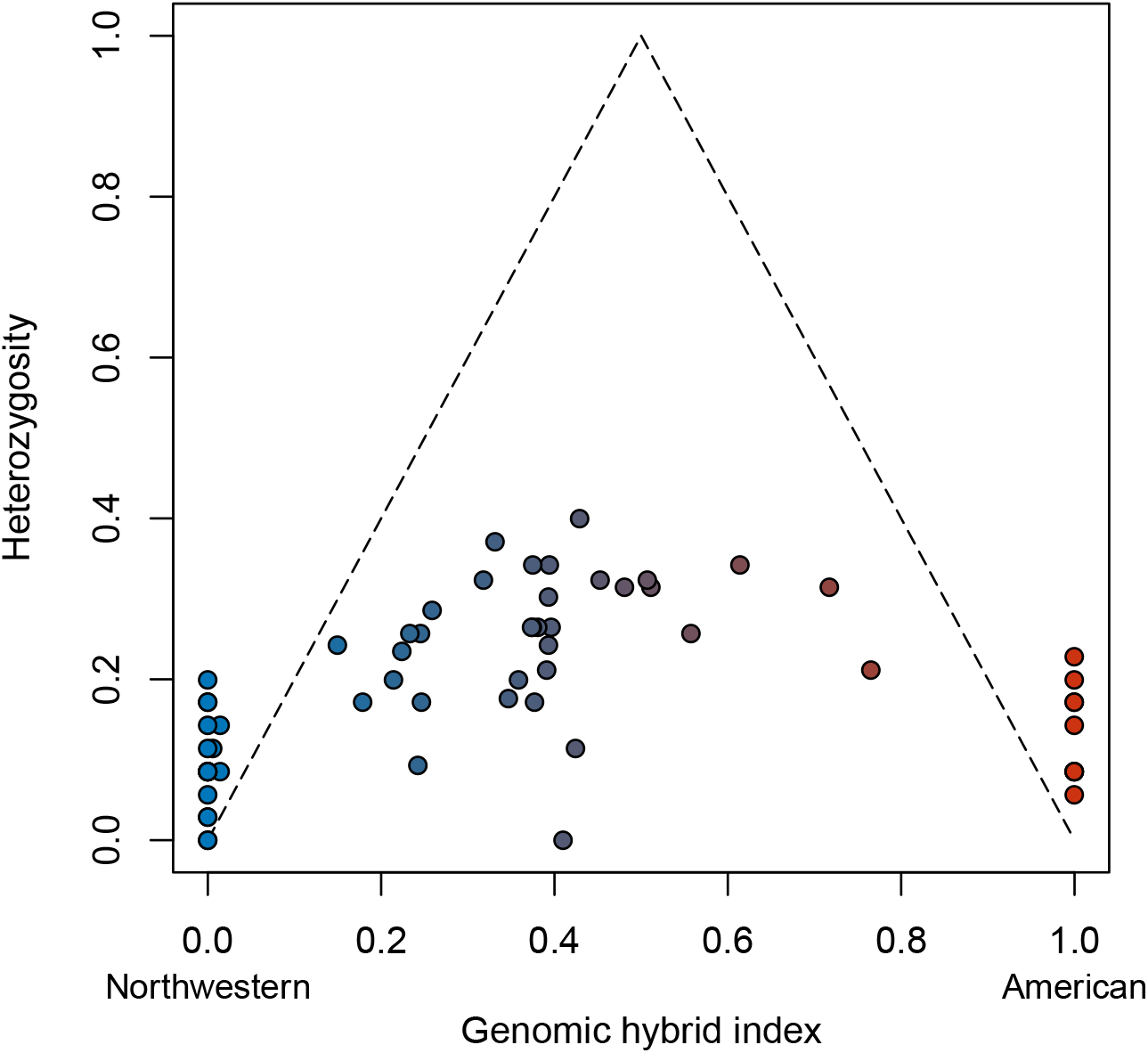
Genomic hybrid index vs. inter-taxon heterozygosity across the Northwestern Crow (blue) and American Crow (red) hybrid zone, based on 34 ancestry-informative nuDNA SNPs (see text). F1 hybrids are expected to have a hybrid index of 0.5 and a heterozygosity of 1.0 for loci fixed in parental individuals. Dotted lines indicate the expected reduction in heterozygosity due to backcrossing (see Methods).

### Nature and extent of mitonuclear concordance and discordance

Nuclear DNA ancestry was generally concordant with mtDNA haplogroup. Crows with pure (>98%) nuDNA ancestry never had a “mismatched” mtDNA haplogroup, but some mitonuclear discordance was evident (Figure 1). Individuals with the American mtDNA haplogroup (n=40) had up to 62% Northwestern nuDNA ancestry (K=2), and crows with the Northwestern mtDNA haplogroup (n=22) had up to 60% American nuDNA ancestry. At six localities, we sampled nuDNA from crows with both American and Northwestern mtDNA haplogroups. Five of these six localities contained only hybrids, and the sixth locality contained two hybrids and two pure Northwesterns.

### mtDNA and nuDNA clines have concordant centers but not widths

We generated Pacific coastal cline models using 218 mtDNA samples from 13 populations (range = 9-31 per population) and nuDNA from 48 individuals in 12 populations (4 per population). The mtDNA and nuDNA clines were both centered in southwestern British Columbia, with overlapping ± 2 LL intervals for cline center. However, the mtDNA cline was over twice as steep as the nuDNA cline (Figure 3), and the ± 2 LL intervals for cline width did not overlap. The mtDNA cline was centered near the latitude of central Vancouver Island (4486 km ± 2 LL range 4313-4620 km) with a width of 1315 km (± 2 LL range 975-1859 km). The nuDNA cline was centered near the latitude of the Strait of Juan de Fuca (4820 km ± 2 LL range 4528-5759 km) with a width of 3563 km (± 2 LL range 2726-5593 km).

**Figure 3:**
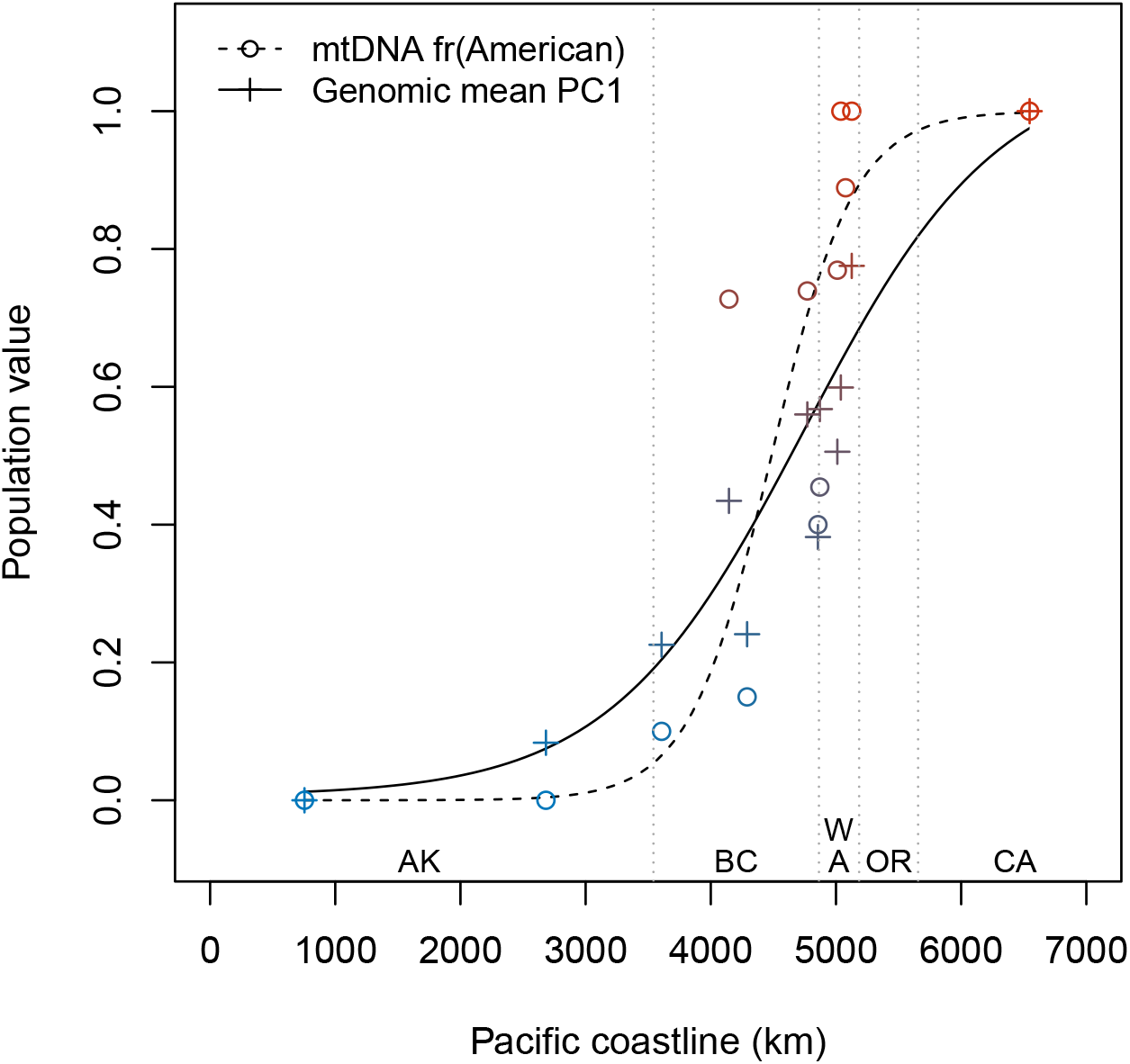
Pacific coastal clines spanning the Northwestern Crow (blue) and American Crow (red) hybrid zone. Circles indicate population frequencies of the American mtDNA haplogroup and crosses indicate population means of nuDNA PC1 (see Methods). Dashed and solid lines represent best-fit cline models (see Methods), and dotted vertical lines indicate state and provincial boundaries.

### Heterogeneous admixture near geographic features of interest

Overall, both mtDNA and nuDNA results indicated a trend of increasing Northwestern ancestry with latitude west of the Cascades and Coast Mountains (Figure 1). However, Vancouver Island crows had a higher frequency of the Northwestern ND2 haplogroup (74%) than crows on the nearby mainland (26%; χ^2^=15.8, p<0.0005). Crows on Vancouver Island also averaged 44% more Northwestern nuDNA ancestry than adjacent mainland crows (74% vs. 51%, t=3.13, df=14, p=0.0074).

Crows within most localities were fairly homogeneous in their ancestry proportions, but localities closest to the Skeena River valley were a notable exception (Figure 1), where within-population ancestry variation was about 9 times higher at these localities than elsewhere (variance parameter = 9.02, likelihood ratio = 85.02, df=1, p=2.9× 10^−20^).

## Discussion

We conducted a population genomic study of American and Northwestern crows to better understand a potential case of speciation without clear morphological differentiation. We found both mtDNA and nuDNA evidence that these crows represent two ancient evolutionary lineages, but we also identified a >1,500 km-wide hybrid zone consisting of a late-generation hybrid swarm. Overall, our results suggest that the American/Northwestern Crow complex represents a compelling example of reticulate evolution (Huson and Bryant 2006, Sessa et al. 2012).

### Potential causes of mitonuclear concordance and discordance

Mitochondrial DNA and nuDNA were concordant with respect to their cline centers, but mtDNA exhibited a steeper cline than nuDNA. This pattern is rare in hybrid zones involving most organisms but is less uncommon in birds (Toews and Brelsford 2012). Even so, only about one-third of avian studies reporting mitonuclear discordance found a mtDNA cline steeper than the corresponding nuDNA cline (Toews and Brelsford 2012). Processes that might explain this pattern include Haldane’s rule, sex-biased asymmetries, and mtDNA selective gradients (Toews and Brelsford 2012).

The narrower mtDNA cline we observed is broadly consistent with Haldane’s rule (Haldane 1922). Haldane’s rule predicts that increased hybrid sterility and reduced hybrid fitness in females, the heterogametic sex in birds, will result in less introgression of maternally inherited mtDNA relative to nuDNA (Carling and Brumfield 2008, McCormack et al. 2011). Patterns consistent with predictions of Haldane’s rule have been reported in other taxa, including butterflies (Devis et al. 1997, Dasmahapatra et al. 2002). To directly test further predictions of Haldane’s rule in American and Northwestern crows, future field studies should look for signs of reduced viability of hybrid females compared to hybrid males (Smith and Rohwer 2000). Although reduced viability and increased sterility in heterogametic hybrids are among the earliest symptoms of reduced hybrid fitness, these disadvantages are less likely to materialize in closely related taxa (Coyne and Orr 1989). Given the Pleistocene divergence of American and Northwestern Crow lineages and the potential for recurrent gene flow during multiple interglacial periods, it is unknown whether sufficient time has passed for such genetic incompatibilities to accumulate. Indeed, the broad hybrid zone we recovered, consisting of late-generation hybrids and backcrosses, suggests that hybrid fitness is not greatly reduced in these crows.

Sex-biased asymmetries such as male-biased dispersal can also lead to a narrow mtDNA cline (Toews and Brelsford 2012), but dispersal in crows is actually biased towards females (Withey and Marzluff 2005). The broad overlap zone of American and Northwestern mtDNA haplogroups also argues against a strong latitudinal selection gradient on mtDNA being a primary driver of the narrower mtDNA cline (e.g., Cheviron and Brumfield 2009). Nonetheless, we cannot rule out that one or more of these latter processes, operating weakly, might be enough to drive the observed difference in the width of mtDNA and nuDNA clines.

### Pleistocene divergence and historical biogeography

When American and Northwestern mtDNA diverged ~381,000 years ago in the late Pleistocene, North America was undergoing extensive glacial advances and retreats at regular ~100,000-year Croll-Milankovich intervals (Muller and MacDonald 1997, Clark et al. 2009). Much of the Pacific Northwest was covered in ice sheets during glacial periods, isolating terrestrial organisms south of the ice sheets or in ice-free northern refugia such as Beringia, Haida Gwaii, or the Alexander Archipelago (Galbreath and Cook 2004, Burg et al. 2006, Anderson et al. 2006, Godbout et al. 2008, Shafer et al. 2010, Geraldes et al. 2018). During interglacial periods, terrestrial organisms expanded from refugial populations into newly ice-free habitats, leading to secondary contact and potentially renewed gene flow between closely related, previously allopatric forms (Shafer et al. 2010). These repeated cycles of isolation and secondary contact created complex and/or reticulate population genetic histories in many of the region’s terrestrial organisms (Hewitt 2004; e.g., Omland et al. 2000, Latch et al. 2009, Kearns et al. 2018).

Northwestern and American crows likely diverged following a similar pattern, with Northwestern Crow populations evolving in isolation in one or more of the ice-free northern refugia while American Crow populations remained south of the ice sheets. Today, Northwestern and American Crow mtDNA haplogroups overlap across most of coastal Washington and all of coastal British Columbia, consistent with post-glacial expansion of previously isolated populations into newly available habitat during one or more interglacial periods. Further sampling of crows on Haida Gwaii, the Alexander Archipelago, and other putative northern refugia might uncover additional genetic diversity with the Northwestern mtDNA haplogroup, which would corroborate the Pleistocene refugia hypothesis (e.g., Krosby and Rohwer 2009, Geraldes et al. 2018).

Our results show that most gene flow between American and Northwestern Crows has occurred on a north-south axis to the west of the Coast Mountains and Cascades. These 2-5 My-old ranges would have restricted east-west gene flow in crows during Pleistocene interglacial periods as they do today (Shafer et al. 2010). However, the elevated Northwestern ancestry we observed in Vancouver Island hybrids indicates that gene flow along the Pacific coastline is not perfectly uniform and may vary based on the geography of islands and water barriers.

Even though geography and genetic data both suggest predominantly north/south gene flow on the Pacific slope of the Coast Mountains and Cascades, we note the potential for limited east-west gene flow across these ranges, especially at major drainages with low passes such as the Fraser and Skeena River valleys. Indeed, the relatively high variation we found in nuDNA ancestry proportions near the Skeena River is consistent with more recent gene flow and backcrossing there compared to other localities. However, despite the higher variance in nuDNA ancestry proportions near the Skeena, the hybrid crows we sampled there all appeared to be late-generation hybrids and backcrosses, like at all other localities we sampled.

### Selection vs. neutral processes and the age of the hybrid zone

A broad hybrid zone consisting of late-generation hybrids and backcrosses is consistent overall with a prolonged period of neutral expansion, but we cannot rule out more recent processes. These crows are human commensals that thrive in disturbed landscapes (Verbeek and Butler 1999, Verbeek and Caffrey 2002), and indigenous peoples have inhabited this hybrid zone for millennia. More recently, European settlers have heavily modified the landscape through deforestation, agriculture, and urbanization. One hypothesis posits that recent land use changes and associated increased habitat heterogeneity may have removed habitat barriers to dispersal that existed before the time of European settlement, increasing opportunities for hybridization between a more maritime Northwestern Crow and a more agrarian American Crow (Marzluff et al. 2005, Haring et al. 2012). Under this scenario, more than a century of crow generations may have been sufficient genetic recombination to dilute highly heterozygous F1s and recent-generation hybrids out of the population.

We note that our ddRAD libraries sampled a diverse yet small fraction of the total nuclear genome. Thus, while our mtDNA and nuDNA results seem mostly consistent with neutral processes and/or Haldane’s rule predominating at the scale of the whole genome, we cannot rule out a role for stronger selection at some loci (e.g., Poelstra et al. 2014).

### Reinterpreting the American/Northwestern Crow complex

Our inaugural population genomic investigation of this long-controversial taxon pair clarifies the status of a species complex for which a lack of diagnostic phenotypic characters has obscured understanding for more than 160 years. In light of our results, past claims of distinct crow species breeding assortatively in sympatry (Brooks 1917, 1942) appear to have been in error, seemingly arising from overzealous or imaginative use of subjective identification criteria. Traditional phenotypic characters for distinguishing American and Northwestern crows included size, ecology, and voice, but these were always controversial when subjected to scrutiny. In the hindsight of our genomic study, it is easier to see why these characters were unreliable. Historically, Northwestern Crows were considered to be diagnostically smaller than American Crows (Baird 1858). In actuality, however, size variation in coastal crow populations is clinal, with northern birds averaging smaller, but with great overlap in measurements among individuals, especially near the range boundary (Rhoads 1893, Johnston 1961). Intertidal habitat use, once thought to be a taxonomic character for Northwestern Crow (Baird 1858), might simply reflect these highly intelligent and adaptive crows responding to local food availability (Cooper 1870). Purported vocal differences (Baird 1858, Suckley and Cooper 1860, Brooks 1917, 1942; Hellmayr 1934) do not seem to correlate with size (Rhoads 1893) or habitat (Johnston 1961) near the range boundary, and individual birds have been observed giving typical vocalizations of both taxa (Johnston 1961). Moreover, crows are oscine passerines that learn song (Beecher and Brenowitz 2005), and individual crows can even change vocalizations when joining a new social group (Brown 1985).

The broad hybrid zone we documented corroborates the work of earlier researchers who painstakingly documented a continuous morphological cline along the Pacific Northwest coast (Rhoads 1893, Johnston 1961), often working against the conventional wisdom of their day. Various authorities over the years have been wildly inconsistent regarding the southern range limit of the Northwestern Crow, placing it anywhere from California (e.g., American Ornithologists’ Union 1895) to Oregon (e.g., American Ornithologists’ Union 1983) to Washington State (e.g., Ridgway 1904, Verbeek and Butler 1999, Verbeek and Caffrey 2002, Clements et al. 2017). These difficulties in identifying a discrete range boundary now make sense given the existence of a broad genomic cline. Notably in our study, however, both mtDNA and nuDNA analyses placed the center of the hybrid zone in southwestern British Columbia, farther north than previous hypotheses based on traditional phenotypic characters (e.g., American Ornithologists’ Union 1998).

The lack of geographic-genetic structuring within American Crow mtDNA (Figure S1) was somewhat surprising given that American Crows are widespread and morphologically variable across their North American distribution (Ridgway 1904, Johnston 1961). However, widespread migration in American Crows (Verbeek and Caffrey 2002, Townsend et al. 2018) combined with occasional long-distance female dispersal (McGowan 2001, Withey and Marzluff 2005) provide a ready ecological mechanism for homogenizing gene flow.

### Prevalence and discovery of cryptic hybrid zones

We leveraged tissue samples from natural history museum specimens to conduct the first population genomic analysis of this morphologically conserved and taxonomically controversial species complex, revealing the existence of a broad, cryptic hybrid zone. This avian hybrid zone in North America involves a well-known taxonomic group in an intensively studied geographic region, so the fact that it remained enigmatic for so long suggests that the global frequency of morphologically cryptic hybrid zones is underestimated. In the case of these crows, a > 160-year taxonomic controversy based on subjective and variable phenotypic characters both motivated our genomic study and portended the existence of a cryptic hybrid zone. We expect that comprehensive population genomic surveys of other morphologically austere taxa will reveal additional cryptic hybrid zones. Furthermore, we encourage researchers to carefully characterize such hybrid zones to facilitate comparisons and synthesis of speciation and hybridization in both morphologically conserved and morphologically distinctive organisms.

## Supporting information

## Acknowledgments

Sharon Birks, Victoria Bowes, Matthew Cleland, Sergei Drovetski, Allen Furnell, Colleen Handel, John Marzluff, Lisa Pajot, and Tracy Sutherland helped salvage or collect crow samples. Robert W. Bryson, Jr., Ross Furbush, Jared Grummer, Adam Leache, Hollie Walsh, and Robert Zink assisted with lab work or bioinformatics. CJ Battey, Cooper French, Laura Frost, Ethan Linck, Sabrina McNew, Lindsey Nietmann, and Yue Shi provided comments or discussion that helped improve the manuscript. Sharon Birks arranged loans from the Genetic Resources Collection at the University of Washington Burke Museum, which provided most of the tissue samples for this study. Additional tissues were provided by the American Museum of Natural History, the Museum of Vertebrate Zoology at Berkeley, the University of Michigan Museum of Zoology, and the Louisiana State University Museum of Natural Science. Any use of trade, firm, or product names is for descriptive purposes only and does not imply endorsement by the U.S. government.

## Data Accessibility

Input files and scripts for running analyses and producing figures are available on GitHub at *https://github.com/slager/crow_hybrid_zone* and have been provided to peer reviewers in a zipped archive. Final scripts and files will be publicly archived (e.g., on Dryad) upon publication. After peer review and prior to final publication, mtDNA sequences will be deposited in GenBank and demultiplexed/filtered nuDNA reads will be archived at the NCBI Short Read Archive or Dryad.

## Author Contributions

DLS, RRH, SR, and JK conceived the study; DLS and JK designed the study; SR, CW, and CVH collected field samples; DLS and KLE conducted the lab work. DLS analyzed the data, interpreted the data, wrote the manuscript, and revised the manuscript with input from all co-authors.

---

Table S1. Detailed sampling information (provided as Supplemental File). Asterisks indicate *Corvus corone*; all other samples are *Corvus brachyrhynchos* or *Corvus caurinus.* Museum abbreviations: UWBM = University of Washington Burke Museum, AMNH = American Museum of Natural History, LSUMZ = Louisiana State University Museum of Zoology, MVZ = Museum of Vertebrate Zoology at Berkeley, UMMZ = University of Michigan Museum of Zoology.

Figure S1. Median-joining haplotype network of mtDNA ND2 in Northwestern and American crows (n=258) with *Corvus corone* outgroup (provided as Supplemental File; see Table S1).

Figure S2. Full map of sampling localities in North America (provided as Supplemental File; see Table S1). The shaded area shows the combined range of American and Northwestern crows. Pie charts show proportion of American (red) and Northwestern (blue) mtDNA ND2 haplogroups at each locality, and total pie area is proportional to mtDNA ND2 sample size.

