## Supplementary figures and images for "Cryptic and extensive hybridization between ancient lineages of American crows"

### Supplementary file 2

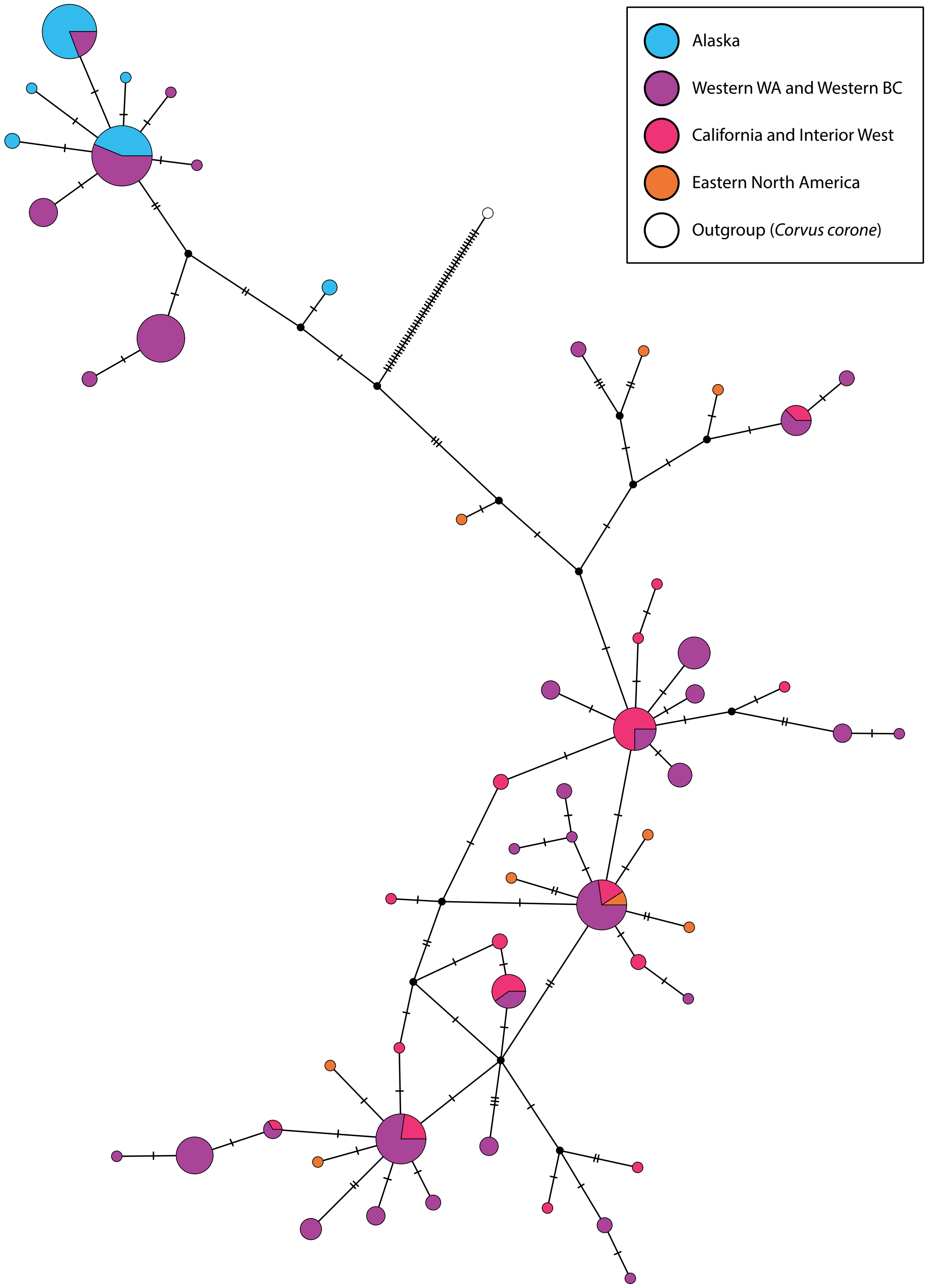

### Supplementary file 3

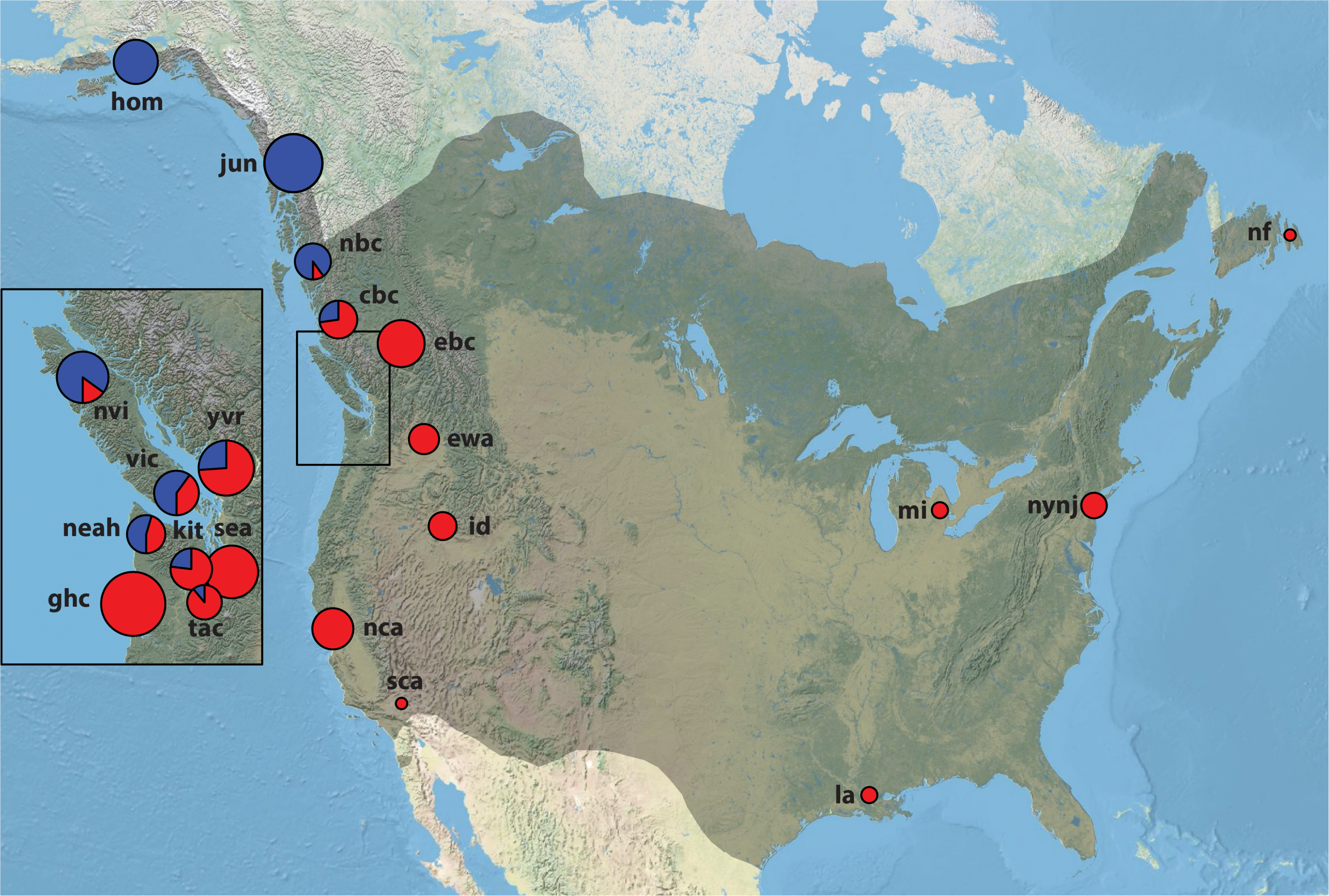
